# *Galleria mellonella* as a novel invertebrate model for studying *Ehrlichia ruminantium* pathogenesis and host-pathogen interactions

**DOI:** 10.64898/2026.04.29.721731

**Authors:** Maëlle Bayet, Christina Nielsen-Leroux, Valérie Rodrigues, Damien F. Meyer

**Affiliations:** CIRAD, UMR ASTRE, Centre for Research and Surveillance on Vector-borne Diseases in the Caribbean, WOAH Reference Laboratory for Heartwater, F-97170 Petit-Bourg, Guadeloupe, France; ASTRE, CIRAD, INRAE, Université Montpellier, Montpellier, France; INRAE, Micalis & AgroParisTech-Université Paris Saclay, Jouy-en-Josas, France

**Keywords:** Ehrlichia ruminantium, Heartwater, Galleria mellonella, Greater wax moth, Invertebrate infection model, Pathogenesis, Host-pathogen interactions

## Abstract

*Ehrlichia ruminantium*, the causative agent of heartwater disease, is an obligate intracellular bacterium that poses significant economic threats to livestock production in endemic regions. Current research models present substantial ethical, logistical, and economic constraints, particularly for studying host-pathogen interactions within arthropod vectors. Here we establish *Galleria mellonella* larvae as a tractable invertebrate infection model for *Ehrlichia ruminantium*, enabling experimental investigation of pathogen persistence and host-pathogen interactions in an arthropod system. Following infection, *G. mellonella* proved susceptible to *E. ruminantium* with moderate mortality and remarkable bacterial persistence. Using rhodamine-labeled bacteria and fluorescence microscopy, we tracked bacterial dissemination from injection sites to systemic distribution in characteristic segmental patterns throughout the larval body. Critically, we confirmed intracellular localization of *E. ruminantium* within hemocytes, the primary immune cells of *G. mellonella*. Quantitative PCR analysis revealed stable bacterial loads over the study period, indicating bacterial persistence within the host.

These findings demonstrate that *E. ruminantium* can hijack the innate immune system of *G. mellonella*, similar to its behavior in natural hosts. The segmental bacterial distribution suggests exploitation of hemolymph circulation and sessile hemocyte populations, providing new insights into potential mechanisms of pathogen persistence. This model offers significant advantages: ethical acceptability, cost-effectiveness, experimental tractability, and compatibility with high-throughput screening approaches. The *G. mellonella* system represents a valuable complement to existing mammalian models and provides a unique platform for investigating arthropod-specific aspects of *E. ruminantium* biology, screening antimicrobial compounds, and understanding mechanisms of immune evasion that may inform strategies for heartwater disease control.

## INTRODUCTION

*Ehrlichia ruminantium* is a Gram-negative, obligate intracellular bacterium belonging to the family *Anaplasmataceae* (order Rickettsiales) that causes heartwater disease, one of the most economically important tick-borne diseases affecting ruminants worldwide (Thomas, 2016; Rodrigues, 2024). This severe and often fatal disease affects domestic and wild ruminants including cattle, sheep, goats, deer, giraffes, and camelids across sub-Saharan Africa, the Indian Ocean region, and the Caribbean (Guadeloupe and Antigua), with ongoing threats of expansion to the American mainland (Allsopp, 2015; Marcelino et al., 2016; Dufleit et al., 2026).

The pathogen exhibits a complex biphasic developmental cycle, alternating between infectious extracellular elementary bodies and intracellular reticulate bodies (Meyer et al., 2023). Transmission occurs primarily through *Amblyomma* ticks during blood feeding, after which *E. ruminantium* replicates in regional lymph nodes before disseminating via the bloodstream to vascular endothelial cells (Rodrigues, 2024). Clinical manifestations include neurological disorders, respiratory distress, hyperesthesia, and the pathognomonic hydropericardium that gives the disease its name (Allsopp, 2015; World Organisation for Animal Health, 2018).

Despite significant economic impact on livestock production, fundamental aspects of *E. ruminantium* pathogenesis, immune evasion mechanisms, and host-pathogen interactions remain poorly understood, particularly within arthropod vectors (Moumène and Meyer, 2016). Traditional research approaches have relied predominantly on mammalian models, including laboratory mice and natural hosts (Seakamela et al., 2024; Simbi et al., 2006; Brayton et al., 2003). While these models have provided valuable insights, they present substantial limitations including ethical considerations, high costs, logistical complexity, and restricted applicability to arthropod-specific research questions.

The need for alternative experimental models is particularly acute given the limited understanding of *E. ruminantium* biology within tick vectors, where the pathogen must survive and potentially replicate before transmission to vertebrate hosts. Recent advances in alternative model systems have highlighted the utility of invertebrate models for studying intracellular bacterial pathogens (Asai et al., 2023; Villani et al., 2025).

*Galleria mellonella*, the greater wax moth larva, has emerged as a powerful, versatile, and cost-effective model organism for studying host-pathogen interactions (Ménard et al., 2021; Villani et al., 2025). This invertebrate system offers several compelling advantages: (i) a functional innate immune system with remarkable similarities to vertebrate immunity, including cellular and humoral responses; (ii) tolerance to mammalian physiological temperatures (37°C and higher), enabling optimal expression of bacterial virulence factors; (iii) ease of handling and manipulation; (iv) ethical acceptability and cost-effectiveness; and (v) compatibility with high-throughput experimental approaches (Kavanagh and Reeves, 2004; Ménard et al., 2021).

*G. mellonella* has been successfully employed to investigate diverse intracellular bacterial pathogens, including *Legionella pneumophila, Coxiella burnetii*, and *Brucella* species, yielding important insights into immune evasion strategies and intracellular persistence mechanisms (Harding et al., 2013; Norville et al., 2014; Barquero-Calvo et al., 2007; Elizalde-Bielsa et al., 2023). These studies have demonstrated the model’s capacity to recapitulate key aspects of pathogen-host interactions observed in mammalian systems while providing unique insights into arthropod-specific immune responses.

To our knowledge, no previous studies have investigated *E. ruminantium* infection in *G. mellonella* or established this system as a model for heartwater research. As summarized in figure 1, this study aims to: (i) establish *G. mellonella* as a viable infection model for *E. ruminantium*; (ii) characterize bacterial localization, dissemination, and persistence within the larval host; (iii) investigate host immune responses and bacterial immune evasion strategies; and (iv) evaluate the model’s potential for advancing our understanding of *E. ruminantium* biology and developing new therapeutic interventions.

**Figure 1.**
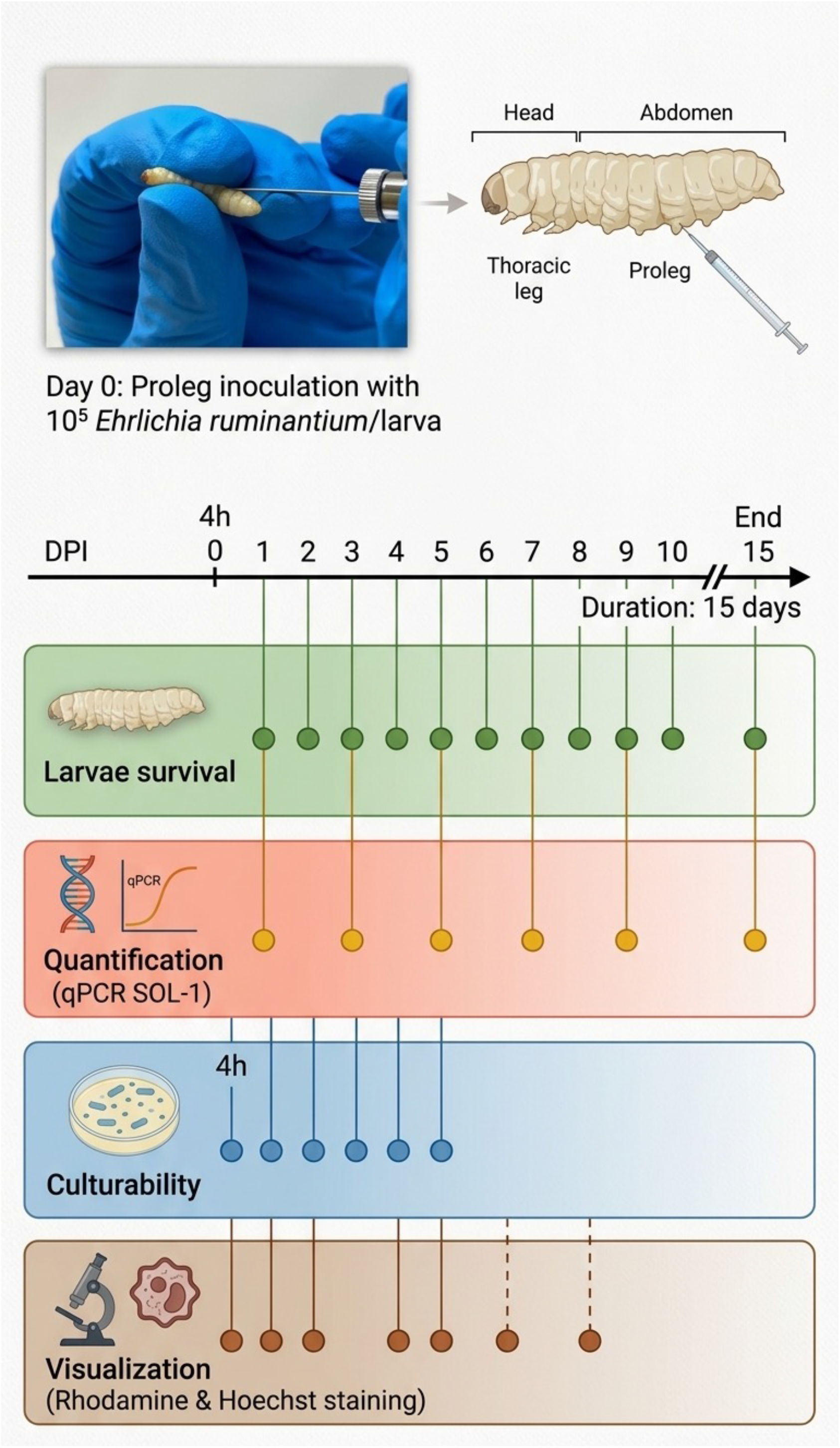
Experimental timeline for the 15-day longitudinal study of *Galleria mellonella* infection with *Ehrlichia ruminantium*. Larvae were inoculated via proleg injection with 10^5^ *E. ruminantium* elementary bodies per larva at day 0. Survival assessment was conducted at fifteen time points throughout the study period. Samples for nucleic acid isolation and qPCR quantification were collected at six time points (days 1, 3, 5, 7, 9 and 15 post-infection). Bacterial cultivability was assessed at six time points (4 hours, and days 1, 2, 3, 4, and 5 post-infection). Visualization experiments using rhodamine and Hoechst staining were performed at seven time points (4 hours, and days 1, 2, 4, 5, 6 and 8 post-infection). DPI, days post-infection.

## METHODS

### Bacterial strains and culture conditions

Bovine aortic endothelial cells (BAEC) were maintained in High Glucose Dulbecco’s modified Eagle’s medium (Gibco, ThermoFisher) supplemented with 10% fetal bovine serum (Gibco, ThermoFisher), 2 mM L-glutamine (Gibco, ThermoFisher), 100 IU/mL penicillin, and 100 μg/mL streptomycin (LifeTechnologies, ThermoFisher). Cells were cultured in 25 cm^2^ tissue culture flasks as monolayers at 37°C in a humidified atmosphere containing 5% CO_2_. BAEC were used up to passage 40 to maintain consistency.

Synchronous *E. ruminantium* infection was established using bacterial suspensions (80-90% infectivity) harvested at 120 hours post-infection (HPI) by mechanical disruption of infected cells through 18G and 26G needles, followed by reinfection at a 1:20 (v/v) ratio. Culture medium was completely replaced at 24 HPI, with half-medium changes performed at 96 HPI to maintain optimal growth conditions.

### *G. mellonella* infection and survival analysis

Elementary bodies were purified by differential centrifugation: initial centrifugation at 300 ×g for 7 min to remove cellular debris, followed by bacterial concentration at 20,000 ×g for 20 min. Bacterial pellets were resuspended in sterile phosphate-buffered saline (PBS), and enumeration was performed using flow cytometry (Accuri C6 Plus, BD Biosciences).

*G. mellonella* last instar larvae (200-250 mg weight) were obtained from MICALIS (INRAE, Jouy en Josas, France) and maintained on bees wax and pollen, at room temperature, until use.

Groups of 10 larvae were injected with 10 μL containing 10^5^ purified elementary bodies per larva into the uppermost right proleg using a sterile Hamilton syringe. Larvae were incubated at 37°C in darkness without food, with survival monitored at 24-hour intervals for 15 days. Death was determined by absence of movement response to gentle mechanical stimulation. Appropriate controls included PBS-injected and untreated groups. All experiments were carried out in triplicate (n=30).

### Bacterial visualization and tracking

For *in vivo* bacterial visualization, *Ehrlichia ruminantium* elementary bodies were labeled with rhodamine B-conjugated polymyxin B (Sigma-Aldrich) at 1.1 μg/mL final concentration, incubated for 30 minutes in darkness, and washed with PBS. Larvae were inoculated with 10^5^ labeled bacteria and observed from 4 HPI to 120 HPI using stereomicroscopy (M205FA, Leica). Individual larvae were immobilized on ice for 10 minutes before observation to prevent movement artifacts.

### Hemocyte analysis and intracellular localization

For hemocyte analysis, larvae were infected with 10^5^ rhodamine-labeled bacteria as described above. Ten microliters hemolymph was collected by aseptically severing the bottom 2 mm of larvae and draining fluid into sterile microcentrifuge tubes. Samples were stained with 100 μL of 10 μg/mL Hoechst 33342 (Sigma-Aldrich) for 15-30 minutes at room temperature before microscopic examination using a fluorescent cell imager (ZOE, BioRad). Observations were done directly on hemolymp without any anticoagulation medium and were conducted from 4 HPI to 120 HPI to track bacterial persistence. The selected images are representative of triplicated experiments.

### Bacterial viability assessment

To confirm viability and infectivity of *E. ruminantium* during larval infection, 10 uL of hemolymph was collected from infected *G. mellonella* and directly inoculated onto BAEC monolayers in T25 flasks. Cultures were maintained under standard conditions with regular medium changes, and bacterial viability was assessed by monitoring morula formation and lysis synchrony characteristic of *E. ruminantium* infection.

### Bacterial quantification by qPCR

Bacterial loads onto larvae were quantified using real-time PCR targeting the pCS20 region (SOL-1). Relative quantification was calculated using the 2^(-ΔΔCt) method with eukaryotic 18S rRNA as the reference gene (total DNA normalization). TaqMan probes were labeled with FAM reporter dye (5’) and BHQ-1 quencher (3’). Primer and probe sequences are provided in Table 1.

**Table 1.**
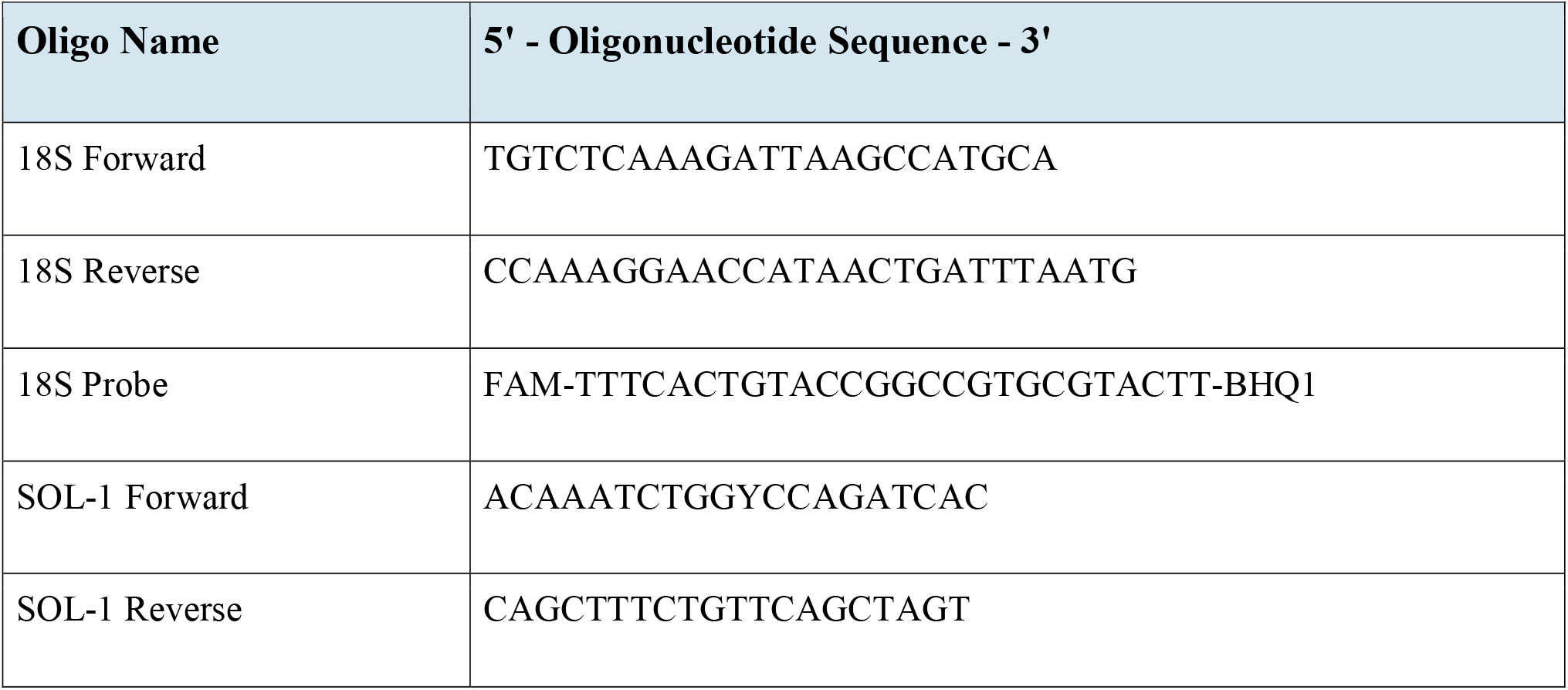
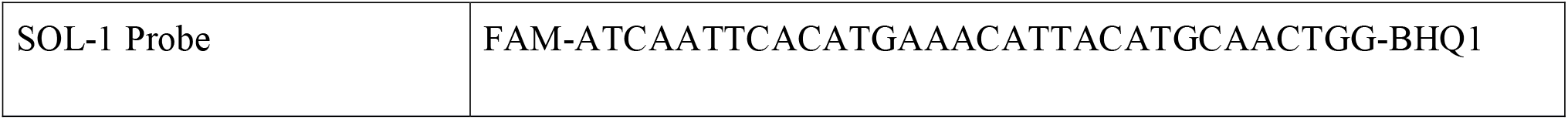
Primers and probes used for qPCR quantification of *E. ruminantium*.

Nucleic acid extraction was performed using the IDEAL-32™ system with ID Gene® Mag Fast Extraction Kit (ID Solutions). Individual larvae were surface-sterilized with ethanol, placed in 2 mL SafeLock tubes (Eppendorf) with 3 mm steel beads, and homogenized using TissueLyser II (Qiagen) at 30 Hz for 2 minutes. Following addition of 340 μL LysFast reagent and 2-minute vortexing, samples were processed according to manufacturer specifications.

Real-time PCR reactions contained 2 μL template DNA, forward primer (250 nM), reverse primer (250 nM), probe (200 nM), and TaqMan Universal PCR Master Mix (ThermoFisher). Cycling conditions comprised initial denaturation at 50°C for 2 min, followed by 95°C for 10 min, then 40 cycles of 95°C for 15 s and 55°C for 1 min.

### Statistical analysis

Statistical analyses were performed using GraphPad Prism version 10 (GraphPad Software, San Diego, CA, USA). Kaplan-Meier survival curves were analyzed using the log-rank (Mantel-Cox) test. Quantitative PCR data were analyzed using one-way ANOVA with Tukey’s multiple comparison test. Results were considered statistically significant at P < 0.05.

## RESULTS

### *G. mellonella* demonstrates susceptibility to *E. ruminantium* infection with characteristic delayed mortality

To establish *G. mellonella* as an infection model for *E. ruminantium*, larvae were challenged with the Gardel strain at 10^5^ bacteria per larva and monitored for 15 days post-infection (Figure 1). Infected larvae exhibited a delayed and moderate mortality, with a final survival rate of 50%, whereas PBS-injected controls showed 100% survival (P = 0.047, log-rank test) (Figure 2A). Monitoring was discontinued after the second death in the control group, and individuals that pupating during the experiment were excluded from the survival analysis. This mortality pattern contrasted markedly with the rapid, high-level mortality typically observed with highly pathogenic bacteria such as *Klebsiella pneumoniae*, suggesting that *E. ruminantium* employs immune evasion strategies that permit prolonged survival while establishing persistent infection. Macroscopic examination revealed progressive changes in infected larvae, including reduced motility and size reduction over the infection period (Figure 2B), while control groups remained healthy throughout the study. Importantly, infected larvae did not exhibit the characteristic melanization response (color change from cream to dark grey/black) typically associated with immune activation against bacterial pathogens. The absence of melanization suggests that *E. ruminantium* may evade or limit activation of the prophenoloxidase pathway, a key component of insect innate immunity.

**Figure 2.**
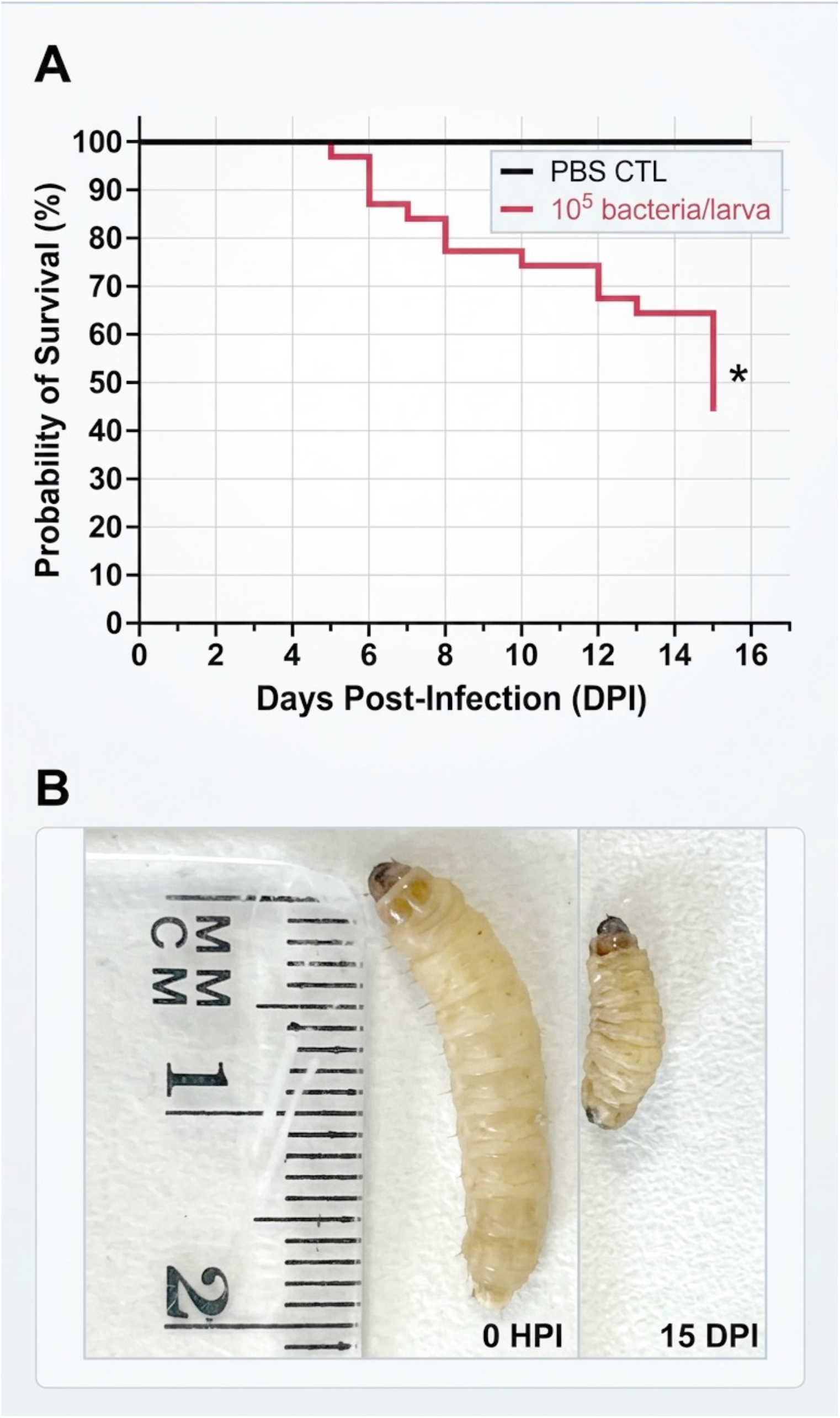
*E. ruminantium* induces delayed mortality in *G. mellonella* with characteristic macroscopic changes. (A) Kaplan-Meier survival curves comparing larvae (n=10 per group) inoculated with 10^5^ *E. ruminantium* bacteria per larva versus PBS-injected controls. Data represent means of three independent replicates. Statistical significance was determined using the log-rank Mantel-Cox test (*P < 0.05). (B) Representative macroscopic changes in *G. mellonella* larvae at 0 and 15 days post-infection, showing reduced size and altered morphology in infected larvae compared to healthy controls. Scale bar, 5 mm.

### E. ruminantium exhibits systematic dissemination and persistent localization within G. mellonella

To track bacterial localization and dissemination, *E. ruminantium* was labeled with rhodamine B-polymyxin B, achieving 97.9% labeling efficiency as confirmed by flow cytometry (Figure 3A). This high labeling efficiency ensured reliable tracking of bacterial distribution throughout the infection period.

**Figure 3.**
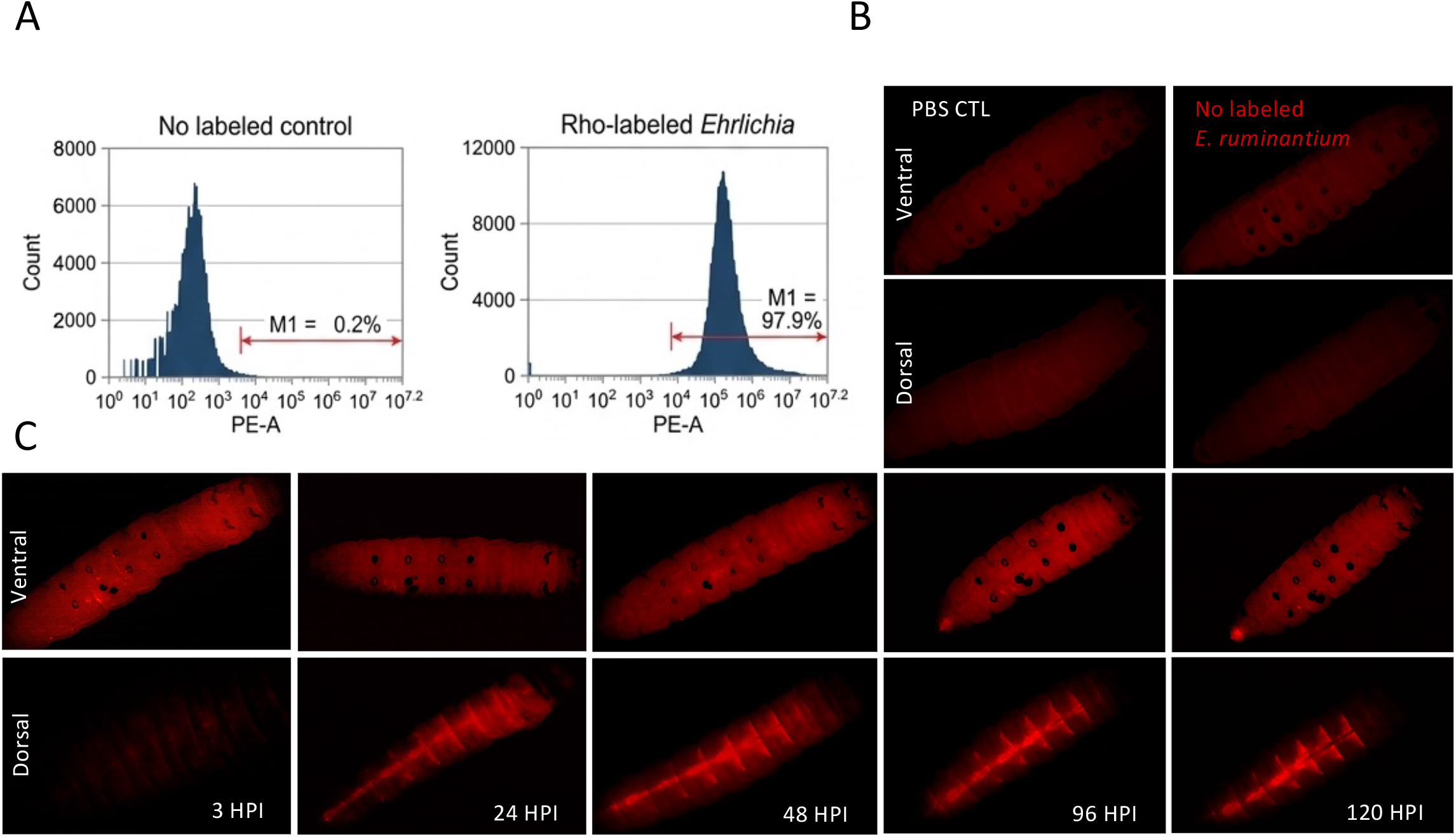
Temporal localization and systemic dissemination of *E. ruminantium* in *G. mellonella*.(A) Flow cytometric validation of rhodamine labeling efficiency. Unlabeled *E.ruminantium* control showing minimal autofluorescence (0.2% positive events). Rhodamine-labeled *E. ruminantium* demonstrating 97.9% labeling with high intensity (high MFI). (B) Control groups showing absence of fluorescence in PBS-injected and unlabeled bacteria-injected larvae. (C) Time-course imaging of bacterial dissemination in larvae infected with rhodamine-labeled *E. ruminantium* (10^7^ bacteria/mL). Individual larvae were tracked by stereomicroscopy from 3 to 120 hours post-infection, showing progressive bacterial spread from the injection site to systemic segmental distribution throughout the larval body. Note the characteristic band-like pattern suggesting compartmentalized bacterial accumulation. HPI, hours post-infection.

Following injection of 10^5^ labeled bacteria into the proleg, fluorescence microscopy revealed dynamic bacterial dissemination patterns (Figure 3B, C). Initially, fluorescence was localized exclusively at the injection site (3 HPI). By 24 HPI, bacterial fluorescence appeared in the dorsal region, indicating successful systemic dissemination. Remarkably, fluorescence distribution exhibited a characteristic repetitive, band-like pattern suggestive of compartmentalized bacterial accumulation associated with larval segmentation. This segmental distribution intensified progressively through 96-120 HPI, with additional bacterial accumulation observed near the rectal region, potentially indicating bacterial trafficking toward elimination routes.

The segmental distribution pattern strongly suggests bacterial exploitation of *G. mellonella* physiological systems, including hemolymph circulation, the tracheal network, or sessile hematopoietic tissues. This systematic distribution was detectable in 100% of infected larvae through 5 days post-infection, demonstrating consistent bacterial establishment and persistence.

### *E. ruminantium* establishes intracellular infection within *G. mellonella* hemocytes

Hemocyte analysis revealed successful intracellular localization of *E. ruminantium* within the primary immune cells of *G. mellonella* (Figure 4). Fluorescence microscopy of hemolymph samples showed normal hemocyte morphology with clear nuclear staining (Hoechst 33342) and distinct intracellular red fluorescence (rhodamine-labeled bacteria) in infected samples. Merged images confirmed colocalization of bacteria within hemocytes, while control samples showed no rhodamine signal, confirming specific bacterial association.

**Figure 4.**
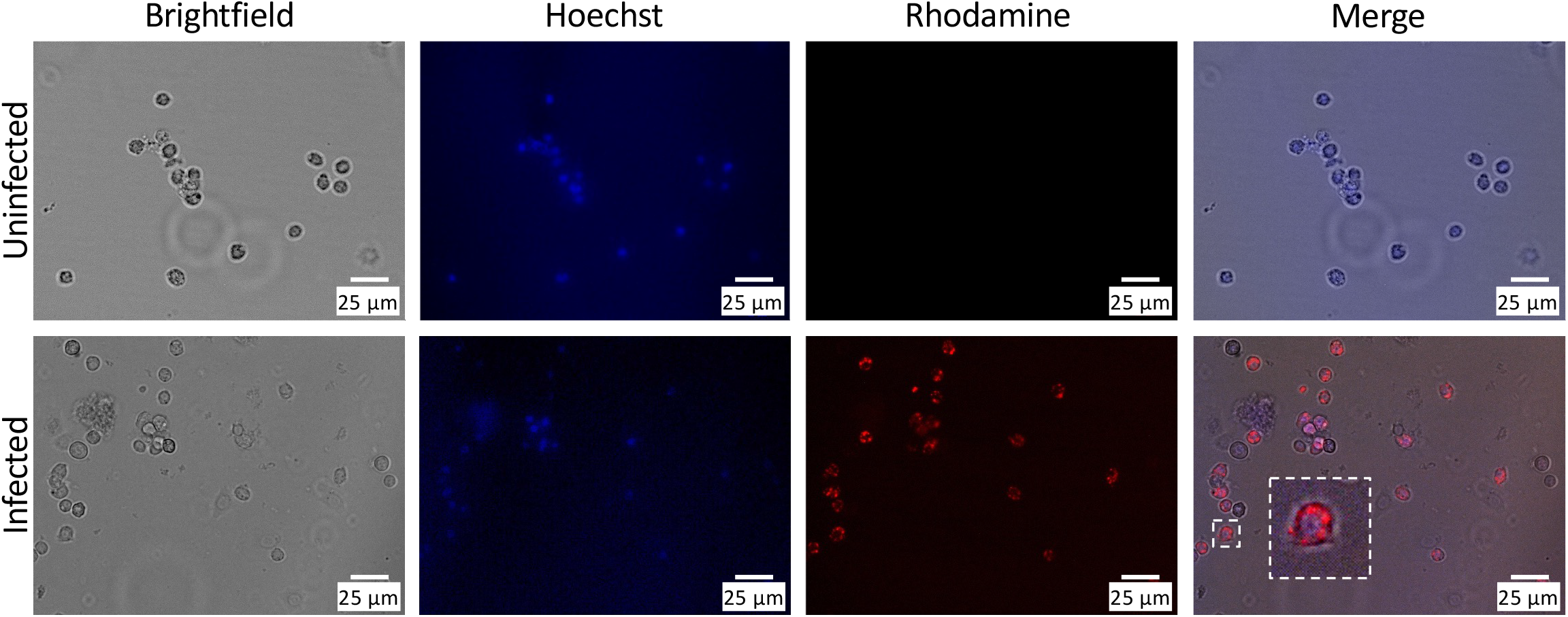
Intracellular localization of *E. ruminantium* within *G. mellonella* hemocytes. Larvae were infected with rhodamine-labeled *E. ruminantium* (10^7^ bacteria/mL), and hemolymph was collected for fluorescence microscopy analysis. Hemocyte nuclei were stained with Hoechst 33342 (blue), and intracellular bacteria appear as rhodamine fluorescence (red). Brightfield images show normal hemocyte morphology in both uninfected and infected samples. The merged images demonstrate clear colocalization of rhodamine-labeled bacteria within hemocytes (white dashed box highlights a representative infected hemocyte). Uninfected controls show no rhodamine signal, confirming specific bacterial association. Scale bar, 25 μm.

Extended analysis demonstrated remarkable bacterial persistence within hemocytes, with intracellular bacteria detectable for at least 6 days post-infection (Figure 5A). Whole-larva imaging confirmed bacterial presence for at least 8 days (Figure 5B), indicating sustained intracellular survival without triggering significant hemocyte destruction or immune clearance.

**Figure 5.**
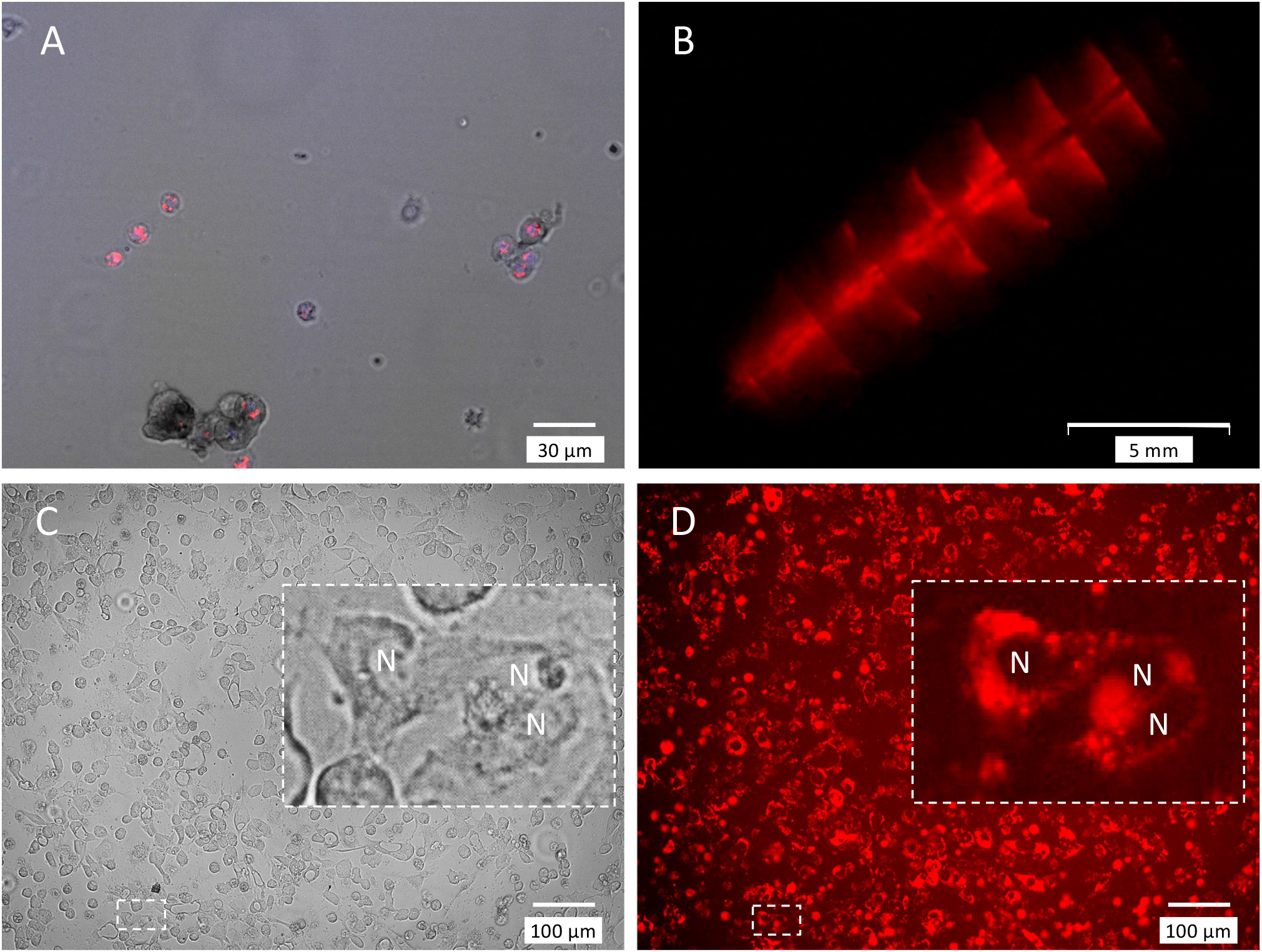
Long-term persistence of viable *E. ruminantium* within *G. mellonella* hemocytes. (A) Fluorescence microscopy of hemolymph collected 6 days post-infection, showing sustained intracellular bacterial presence within hemocytes. Nuclei are stained with Hoechst (blue) and bacteria with rhodamine (red). Scale bar, 30 μm. (B) Stereomicroscopy demonstrating whole-larva bacterial fluorescence persistence at 8 days post-infection. Scale bar, 5 mm. (C) Viability confirmation through re-infection assay. Hemolymph from infected *G. mellonella* was inoculated onto bovine aortic endothelial cell (BAEC) monolayers. At 4 days post-inoculation, characteristic morulae formation, lysis areas, and fluorescent bacteria within BAEC confirm maintenance of bacterial viability and infectivity throughout the larval infection period. Scale bar, 100 μm. N, nuclei; DPI, days post-infection.

### Bacterial viability and infectivity are maintained during *G. mellonella* infection

To confirm that fluorescent signals represented viable, infectious bacteria rather than fluorescent debris, hemolymph from infected larvae was directly inoculated onto BAEC monolayers. Microscopic examination at 4 days post-inoculation revealed characteristic morulae formation and synchronized lysis areas typical of active *E. ruminantium* infection (Figure 5C). Fluorescently labeled bacteria were clearly visible within BAEC, confirming that rhodamine labeling did not impair bacterial viability or infectivity (Figure 5D). This experiment definitively established that *E. ruminantium* maintains full pathogenic potential throughout the *G. mellonella* infection period.

### *E. ruminantium* establishes stable, persistent infection with minimal bacterial clearance

Quantitative PCR analysis of bacterial loads revealed remarkable stability over the 15-day observation period (Figure 6). Bacterial DNA levels remained consistently detectable with only minor fluctuation (approximately 0.5 log variation), indicating that *E. ruminantium* successfully established a stable replicative niche within *G. mellonella* without significant bacterial clearance by host immune mechanisms.

**Figure 6.**
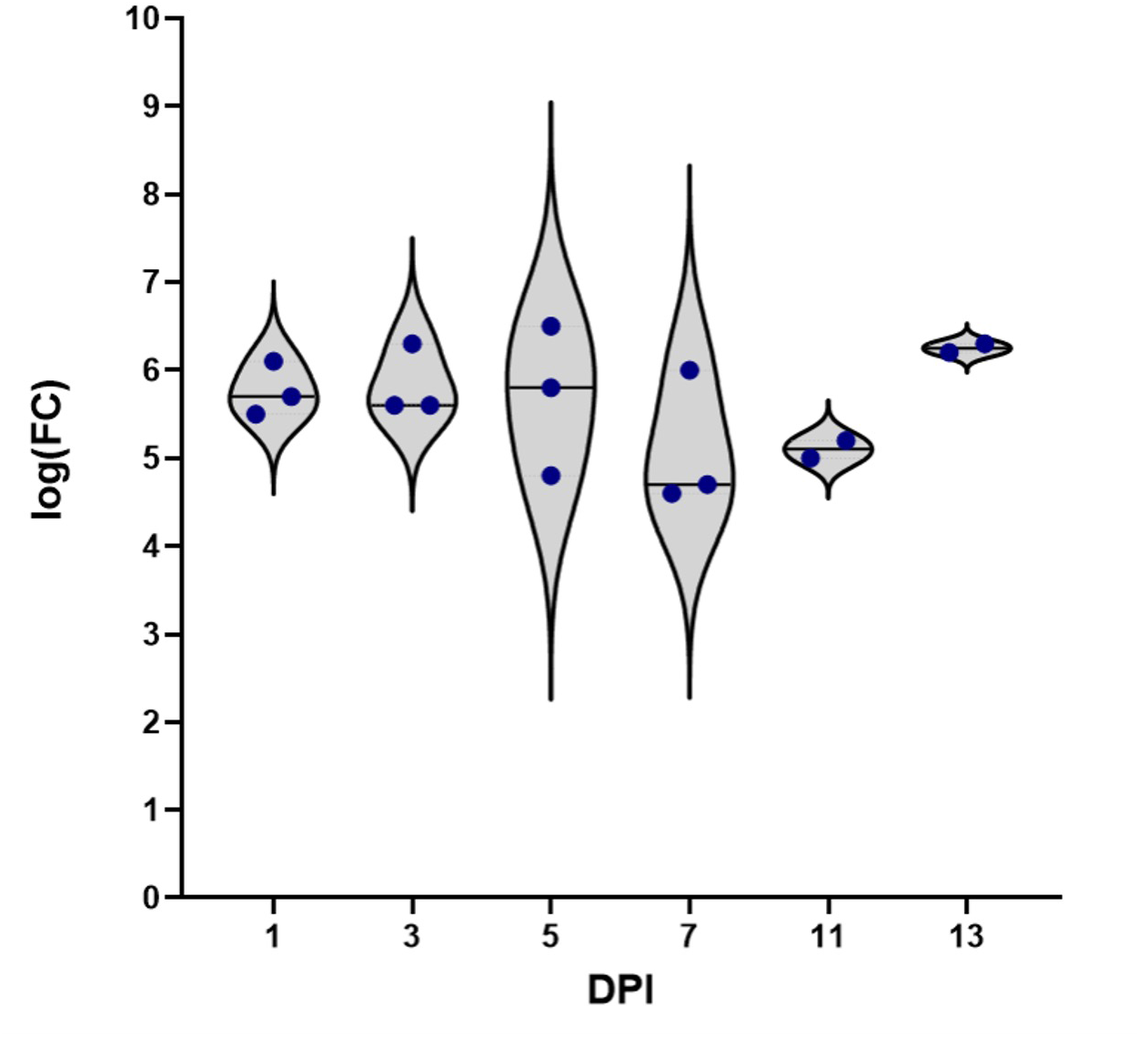
*E. ruminantium* establishes stable, persistent infection in *G. mellonella*. Bacterial loads were quantified by the SOL1 qPCR targeting the pCS20 gene in larvae (n=3) inoculated with 10^5^ bacteria per larva. Samples were collected at indicated time points and bacterial enumeration assessed using the 2^-ΔΔCt^ method with eukaryotic 18S rRNA as reference gene. Violin plots show individual data points and distribution. Bacterial loads remained stable throughout the 15-day observation period with minimal variation (P > 0.05, one-way ANOVA), indicating successful establishment of persistent infection without significant bacterial clearance. DPI, days post-infection.

The stable bacterial loads, combined with sustained viability and minimal mortality, demonstrate that *E. ruminantium* employs sophisticated immune evasion strategies that enable long-term persistence within the invertebrate host. This persistence pattern mirrors the pathogen’s behavior in natural mammalian hosts, where chronic infection and immune modulation are characteristic features of heartwater pathogenesis.

## DISCUSSION

This study represents the first successful establishment of *G. mellonella* as an infection model for *E. ruminantium*, providing a novel platform for investigating host-pathogen interactions in this economically important tick-borne disease. Our findings demonstrate that *E. ruminantium* can effectively infect, persist within, and modulate the immune system of *G. mellonella*, recapitulating key aspects of pathogenesis observed in natural hosts while offering unique insights into invertebrate-pathogen interactions.

### Advantages and applications of the *G. mellonella* model for *E. ruminantium* research

The *G. mellonella* model offers several significant advantages over traditional mammalian systems for *E. ruminantium* research. First, the ethical acceptability and cost-effectiveness of this invertebrate model enable high-throughput experimental approaches that would be prohibitive with mammalian models. Second, the ability to maintain larvae at 37°C ensures optimal expression of bacterial virulence factors that are temperature-regulated and critical for pathogenesis in mammalian hosts. Third, the experimental tractability allows controlled dose administration and precise temporal sampling that facilitates mechanistic studies of infection dynamics.

Perhaps most importantly, this model provides a bridge between mammalian and arthropod research contexts. While *G. mellonella* and *Amblyomma* ticks differ substantially in biology and ecology, both are arthropods with innate immune systems that share fundamental features including cellular immunity, humoral responses, and antimicrobial peptide production (Ribeiro and Brehélin, 2006; Kavanagh and Reeves, 2004). This model therefore offers opportunities to investigate arthropod-specific aspects of *E. ruminantium* biology that cannot be studied in mammalian systems, while providing insights that may be relevant to understanding some pathogen-tick interactions aspects.

### Immune evasion strategies and persistence mechanisms

The delayed, moderate mortality observed in *E. ruminantium*-infected larvae contrasts sharply with the rapid, high-level mortality induced by typical Gram-negative pathogens such as *Klebsiella pneumoniae* (Elizalde-Bielsa et al., 2023). This pattern, combined with the absence of melanization responses, suggests that *E. ruminantium* employs a ‘stealth’ infection strategy similar to that described for *Brucella* species (Barquero-Calvo et al., 2007). The lack of melanization is particularly significant, as this response represents a crucial component of invertebrate immunity involving prophenoloxidase activation and pathogen encapsulation (Cerenius et al., 2008).

Several mechanisms may contribute to *E. ruminantium* immune evasion in *G. mellonella*. The type IV secretion system (T4SS) VirB/D4, which is conserved across *Ehrlichia* species, translocates bacterial effectors into host cell cytoplasm, subverting cellular pathways and immune responses (Rikihisa, 2017; Celli et al., 2003). Studies with *Legionella pneumophila* have demonstrated that T4SS mutants show reduced virulence and impaired intracellular persistence in *G. mellonella* (Harding et al., 2012), suggesting that similar mechanisms may operate in *E. ruminantium* infections.

Additionally, the absence of canonical lipopolysaccharide (LPS), a major pathogen-associated molecular pattern in many Gram-negative bacteria, may contribute to immune evasion by reducing pathogen-associated molecular pattern (PAMP) recognition. In *Ehrlichia* and other members of the *Anaplasmataceae*, the lack of canonical LPS and peptidoglycan is thought to reduce innate immune detection and favor intracellular survival (McBride et al., 2011). In our model, the limited mortality and absence of visible melanization are consistent with this hypothesis, although the underlying mechanisms remain to be determined.

### Bacterial dissemination and cellular targeting strategies

The segmental bacterial distribution pattern observed in infected larvae provides important insights into *E. ruminantium* dissemination strategies. This pattern likely reflects exploitation of hemolymph circulation and the distribution of sessile hemocyte populations attached to tissues and the body wall (King and Hillyer, 2013; Leitão and Sucena, 2015). Sessile hemocytes play crucial roles in controlling microbial infections and are distributed in segmental patterns that could explain the observed bacterial localization.

The successful intracellular localization of *E. ruminantium* within *G. mellonella* hemocytes parallels the pathogen’s behavior in mammalian endothelial cells, where it establishes replicative niches within membrane-bound vacuoles. This cellular targeting strategy appears to be conserved across diverse host species, suggesting fundamental aspects of *E. ruminantium* pathogenesis that transcend phylogenetic boundaries. The ability to establish intracellular infection within primary immune cells represents a particularly sophisticated virulence strategy that enables immune evasion while providing access to nutrients and protection from extracellular immune effectors.

### Future research directions and applications

The established *G. mellonella* model opens numerous avenues for advancing *E. ruminantium* research. Transcriptomic and proteomic analyses of infected larvae could identify host factors influencing susceptibility and bacterial effectors involved in immune modulation. Screening of mutant bacterial strains, particularly those defective in T4SS function, could elucidate specific virulence mechanisms and their contributions to persistence.

The model’s compatibility with high-throughput approaches makes it particularly valuable for antimicrobial compound screening and vaccine development research. The ability to rapidly assess large numbers of potential therapeutic interventions in a cost-effective, ethically acceptable system could accelerate the development of new strategies for heartwater control.

Furthermore, comparative studies examining *E. ruminantium* behavior in *G. mellonella versus* other invertebrate models or arthropod cell cultures could provide insights into conserved *versus* host-specific aspects of pathogenesis. Such studies may ultimately inform our understanding of *E. ruminantium* interactions with natural tick vectors.

### Model limitations and validation considerations

While the *G. mellonella* model offers significant advantages, important limitations must be acknowledged. The substantial evolutionary distance between lepidopteran larvae and natural ruminant or tick hosts necessitates careful validation of findings in established mammalian models or tick systems. Additionally, the simplified immune system of *G. mellonella* lacks adaptive immunity components that may be crucial for certain aspects of heartwater pathogenesis.

The differences in life cycles, feeding habits, and environmental conditions between *G. mellonella* and *Amblyomma* ticks limit direct extrapolation of results to tick biology. However, the model’s value lies not in replacing existing systems but in complementing them by providing accessible experimental approaches for mechanistic studies that would be challenging or impossible in other systems.

## Conclusions

This study establishes *G. mellonella* as the first tractable invertebrate model for *E. ruminantium* infection and reveals intracellular persistence within hemocytes together with a striking segmental dissemination pattern in an arthropod host. The model recapitulates key aspects of pathogenesis while offering unique advantages for mechanistic studies, compound screening, and high-throughput experimental approaches.

The *G. mellonella* system represents a significant addition to the toolkit available for *E. ruminantium* research, offering opportunities to accelerate discovery of new therapeutic targets and interventions for heartwater disease control. While this model should complement rather than replace existing systems, its accessibility and experimental tractability position it as a valuable experimental platform for advancing our understanding of this important veterinary pathogen.

## ACKNOWLEDGMENTS

We are grateful to the MICALIS insect facility for providing *G. mellonella* larvae and technical support. We thank Mélanie Dhune and Naomie Pature for technical assistance. We acknowledge the WOAH Reference Laboratory for Heartwater and the MiVeC biology resource center for providing the *E. ruminantium* Gardel strain and infrastructure support.

